# TGFβ limits Myc-dependent TCR-induced metabolic reprogramming in CD8^+^ T cells

**DOI:** 10.1101/2022.03.30.486376

**Authors:** Helen Carrasco Hope, Gabriella Pickersgill, Pierpaolo Ginefra, Nicola Vannini, Graham P. Cook, Robert J. Salmond

## Abstract

T cell activation is dependent upon the integration of antigenic, costimulatory and cytokine-derived signals and the availability and acquisition of nutrients from the environment. Furthermore, T cell activation is accompanied by reprogramming of cellular metabolism to provide the energy and building blocks for proliferation, differentiation and effector function. Transforming growth factor β (TGFβ) has pleiotropic effects on T cell populations, having both an essential role in the maintenance of immune tolerance but also context-dependent pro-inflammatory functions. We set out to define the mechanisms underpinning the suppressive effects of TGFβ on mouse CD8^+^ T cell activation. RNA-sequencing analysis of TCR-stimulated T cells determined that Myc-regulated genes were highly enriched within gene sets downregulated by TGFβ. Functional analysis demonstrated that TGFβ impeded TCR-induced upregulation of amino acid transporter expression, amino acid uptake and protein synthesis. Furthermore, TCR-induced upregulation of Myc-dependent glycolytic metabolism was substantially inhibited by TGFβ treatment with minimal effects on mitochondrial respiration or mTOR activation. Thus, our data suggest that inhibition of Myc-dependent metabolic reprogramming represents a major mechanism underpinning the suppressive effects of TGFβ on CD8^+^ T cell activation.

## Introduction

CD8^+^ T cell function is determined by the sum of inputs received from the environment, including antigenic and co-stimulatory signals, cytokines, or nutrient availability. In recent years, research focused on describing the metabolic regulation of immunity has defined cell-intrinsic modulatory networks that tightly regulate T cell responses through the integration of these environmental cues. For example, CD8^+^ T cells respond to TCR-stimulation by inducing metabolic reprogramming in part mediated by the activation of the transcription factor Myc (1). Myc activity boosts aerobic glycolysis and the upregulation of amino acid transporters such as SLC7A5, which are crucial to sustain the energetic and biosynthetic demands required for clonal expansion and differentiation (1–3). On the other hand, signals such as CD28 co-stimulation (4), or the cytokines IL-7 and IL-15 (5, 6), have been shown to promote mitochondrial metabolism, key to determine the engagement of memory formation. Moreover, the manipulation of metabolic pathways has proved to be a powerful tool to shape T cell activity and develop therapeutic strategies (7). Thus, studying how diverse signals influence T cell metabolism provides valuable knowledge to further understand regulatory mechanisms of T cell function (or dysfunction) and find therapeutic approaches.

Transforming growth factor β (TGFβ) is a pleiotropic cytokine that acts on both immune and non-immune cells, regulating many biological processes including cell migration, wound healing and angiogenesis. In T lymphocytes, TGFβ is involved in the development and suppressive activity of regulatory T cells as well as in the restriction of effector T cell activation, therefore playing an important function in preventing autoimmunity. Furthermore, TGFβ is a key immunosuppressive cytokine secreted by pro-tumour cells and cancer cells to mediate immune evasion and promote metastasis. Previous work determined that TGFβ inhibits mitochondrial respiration and limits IFNγ secretion of human CD4^+^ T cells through direct interaction of SMAD proteins with the mitochondria (8). Further reports demonstrated that TGFβ restricts mitochondrial metabolism and glycolysis in NK cells by decreasing mammalian target of rapamycin (mTOR) activity (9, 10). It has been more recently described that TGFβ can also limit mTOR activity in CD8^+^ T cells which, paradoxically, promotes mitochondrial metabolism and protects T cells from entering a terminally exhausted phenotype in the context of LCMV-induced chronic stimulation (11). These discrepancies indicate that the roles and mechanisms of TGFβ in the immune response are highly variable depending on cell type and context.

In the current work, we describe how TGFβ modulates early stages of CD8^+^ T cell activation. Transcriptomic analysis identified the transcription factor Myc as a key target of TGFβ signalling. In this regard, TGFβ treatment restricted TCR-induced Myc-dependent metabolic reprogramming. Specifically, TGFβ-treated CD8^+^ T cells display impaired expression of genes involved in glycolytic and amino acid (AA) metabolism which subsequently limited nutrient uptake and appropriate engagement of glycolysis or protein synthesis upon T cell activation. However, TGFβ did not impact upon mitochondrial or mTOR activity. This study sheds light into a mechanism by which TGFβ regulates early CD8^+^ T cell activation and metabolism through inhibition of Myc expression.

## Results

### TGFβ preferentially inhibits CD8^+^ T cell responses to low affinity antigen

To define the inhibitory capacity of TGFβ on CD8^+^ T cell activation, OT-I TCR transgenic T cells were activated with high affinity ovalbumin peptide SIINFEKL (OVA-N4) or low affinity SIITFEKL (OVA-T4) ± TGFβ for 48 hours. Assessment of forward scatter-side scatter parameters by flow cytometry demonstrated that TGFβ treatment limited TCR-induced OT-I T cell growth with a more prominent effect on low affinity OVA-T4-induced responses (Figure 1A). TGFβ inhibited OVA-N4-induced granzyme B expression but had little effect on surface expression levels of either CD25 or CD71 (Figure 1B). By contrast, TGFβ treatment strongly inhibited low affinity OVA-T4-induced CD25, CD71 and granzyme B expression (Figure 1C). As previously reported (12), CD44 expression was elevated following peptide + TGFβ treatment as compared to peptide alone, irrespective of antigen affinity (Figure 1, B-C).

**Figure 1:**
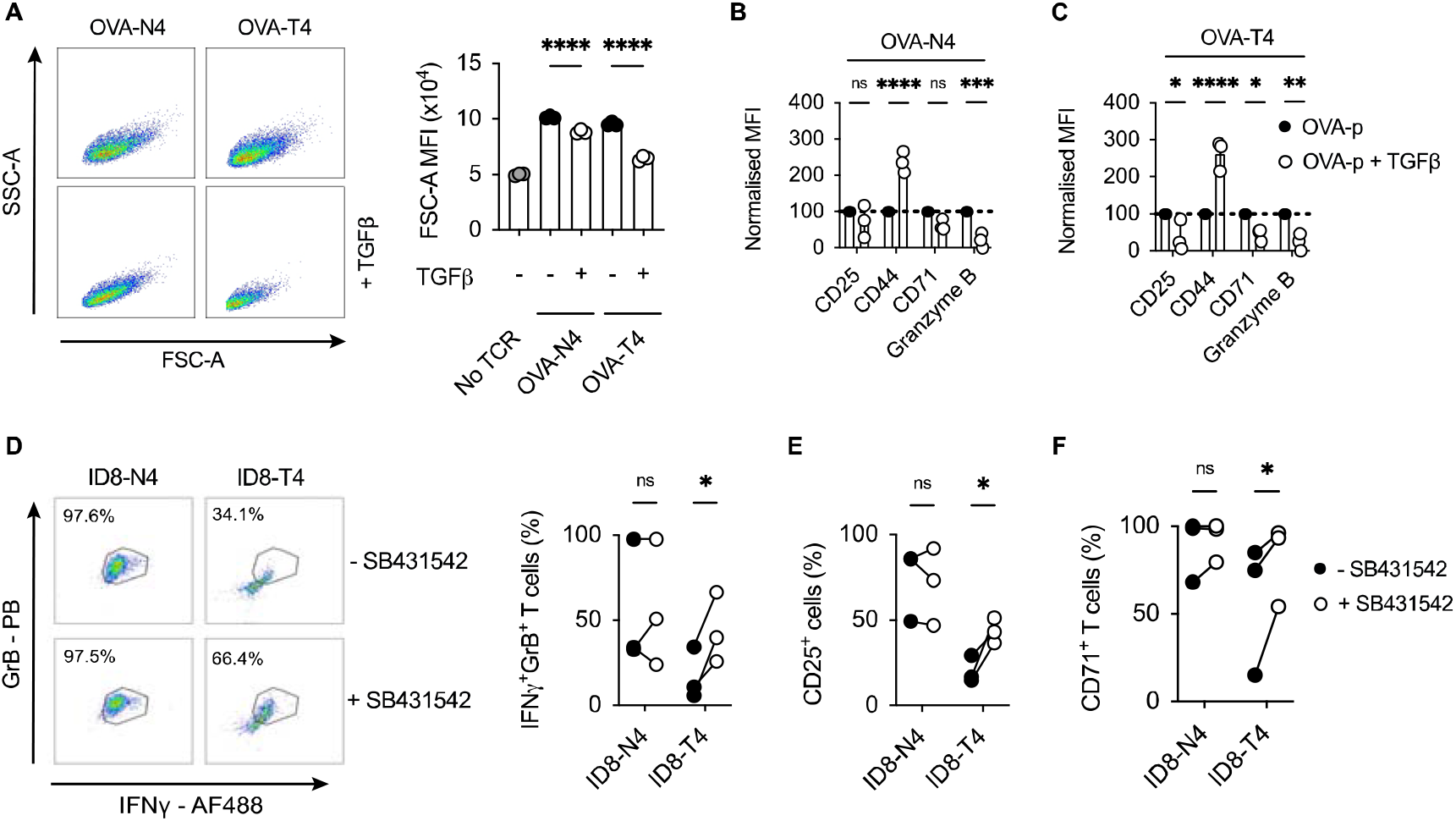
TGFβ inhibits CD8^+^ T cell activation in response to low affinity antigen. **(A-C)** OT-I T cell receptor (TCR) transgenic T cells were stimulated with cognate SIINFEKL (OVA-N4) or low affinity SIITFEKL (OVA-T4) peptides in the presence or absence of TGFβ for 48 hours *in vitro*. **(A)** T cell growth was assessed by analysis of forward and side scatter area (FSC-A/SSC-A) by FACS. **(B, C)** Cell surface expression of activation markers (CD25, CD44, CD71) or intracellular granzyme B were determined by flow cytometry. Data shown represent mean fluorescence intensity (MFI) values for markers, normalised to control (TCR-stimulated) levels. **(D, E)** OT-I T cells were co-cultured in the presence of ID8-OVA-N4 or ID8-OVA-T4 ovarian carcinoma cells in the presence or absence of TGFβRII inhibitor SB431542 for 48 hours *in vitro*. Values in dotplots represent % Granzyme B-IFNγ double-positive cells. **(D)** Data show representative FACS dotplots of intracellular IFNγ and granzyme B expression. Cell surface expression of CD25 (E) and CD71 (F) were determined by flow cytometry. Data points joined by lines represented paired SB431542-treated and untreated samples **(E, F).** Data are from 1 of 3 experiments **(A-D)** or are pooled data from 3 experiments **(E).** Individual data points represent technical replicates **(A-C)** or biological replicates **(D-F)** (n=3). NS – not significant, * p<0.05, ** p<0.01, *** p<0.001, **** p<0.0001 as assessed by 1-way ANOVA, with Tukey’s multiple comparisons test.

Many cancer cell types, including ovarian carcinomas, secrete TGFβ to suppress T cell responses (13–15). We took advantage of mouse ID8 ovarian carcinoma cell lines expressing OVA-peptides to assess the impact of tumour cell-derived TGFβ on OT-I T cell activation. FACS analysis demonstrated that a higher proportion of OT-I T cells co-expressed granzyme B and IFNγ following co-culture for 48h in the presence of ID8 cells expressing OVA-N4 as compared to OVA-T4 (Figure 1D). Addition of the TGFβRII inhibitor SB431542 to OT-I:ID8-OVA-T4 co-cultures doubled the proportions of IFNγ ^+^granzyme B^+^ OT-I cells, but did not impact upon OT-I T cell responses to ID8-OVA-N4 tumour cells. Similarly, OT-I expression of CD25 and CD71 following co-culture with ID8-OVA-T4, but not ID8-OVA-N4 cells, were elevated upon TGFβRII blockade (Figure 1, E-F). Together, these data indicate that both recombinant and tumour cell-derived TGFβ limit TCR-induced T cell activation with a substantially greater effect on responses to lower affinity antigens.

### TGFβ inhibits the TCR-induced Myc-dependent transcriptome

To define the global impact of TGFβ on the TCR-induced transcriptional programme, OT-I CD8^+^ T cells were stimulated for 24h with OVA-T4 ± TGFβ and RNA-Sequencing performed. A total of 5800 transcripts were found to be differentially expressed (adjusted *p* value < 0.05, no fold-change cut-off) between TGFβ-treated and -untreated groups, with 3248 upregulated and 2552 downregulated (Figure 2A). The well-characterized TGFβ target genes *Gzmb* and *Eomes* were amongst the most-downregulated, whilst chemokine receptor-encoding *Ccr8* was amongst the most-upregulated (Figure 2B). Kyoto Encyclopedia of Genes and Genomes (KEGG) pathway analysis of the TGFβ-upregulated gene list (fold-change > 1.5, n=2141) identified “Th17 cell differentiation”, “Leishmaniasis” and “Inflammatory bowel disease” to be among the top upregulated pathways (Supplemental Table 1), consistent with previously described roles for TGFβ in T_h_17/T_c_17/Treg differentiation and function (15, 16).

**Figure 2:**
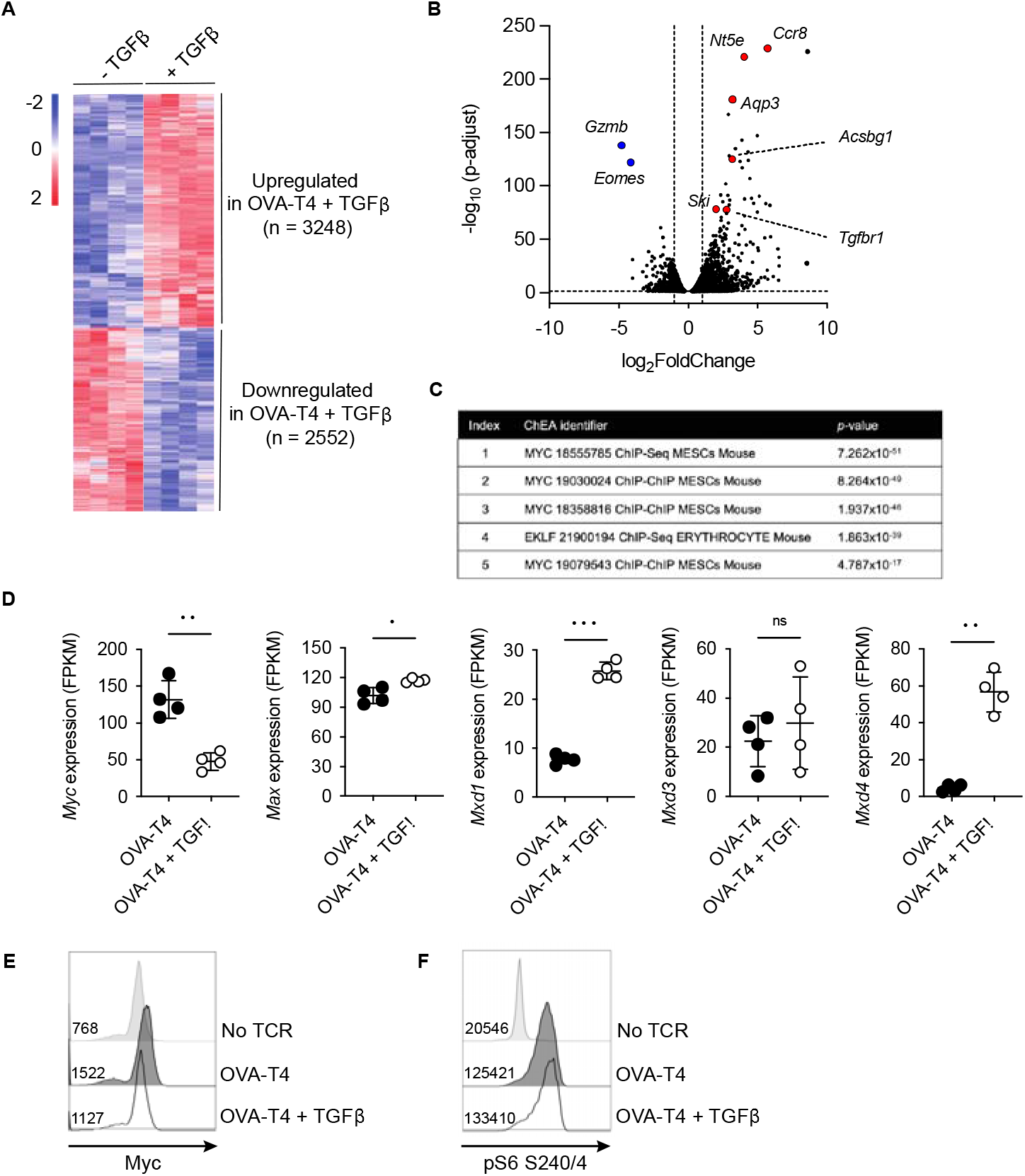
TGFβ impedes the Myc-dependent transcriptome independently of effects on mTOR. OT-I TCR transgenic T cells were stimulated with SIITFEKL (OVA-T4) peptide in the presence or absence of TGFβ for 24 **(A-D, F)** or 48 **(E)** hours. RNA-Sequencing (RNA-Seq) analysis was performed (A) The heatmap shows log_2_ fold changes of differentially expressed genes (p adjust <0.05). (B) Volcano plot of RNA-Seq data identifies genes substantially downregulated (*Gzmb, Eomes*) or upregulated (*Nt5e, Ccr8, Aqp3, Ski, Tgfbr1, Acsbg1*) in TGFβ -treated OT-I T cells. **(C)** ChIP-X enrichment analysis (ChEA) within the Enrichr platform identified Myc-regulated genes as highly enriched within the TGFβ-downregulated gene set. (D) Levels of gene expression of *Myc* and Myc-associated genes. Data shown represent fragments per kilobase million (FPKM) values taken from RNA-Seq analysis. Histograms show levels of Myc expression (E) and phosphorylated ribosomal protein S6 (pS6) (F) as determined by intracellular staining and flow cytometry. In histograms, figures represent MFI values for each condition. Individual data points from RNA-Seq data **(A-D)** represent biological replicates (n=4), whilst FACS plots represent data from 1 of 3 repeated experiments. NS – not significant, * p<0.05, ** p<0.01, *** p<0.001 as determined using DESeq2 package with p-values adjusted using Benjamini-Hochberg procedure.

We focused on analysis of the transcripts downregulated following OVA-T4 + TGFβ treatment (“TGFβ-down”, n = 1139, fold-change > 1.5). Use of the ChIP-X Enrichment Analysis (ChEA) tool (17) within the Enrichr platform (18) identified Myc target genes as highly enriched within the TGFβ-down gene list (Figure 2C). *MYC* was previously identified as a TGFβ-regulated, SMAD-repressed gene in human keratinocytes (19), and to be transcriptionally downregulated in TGFβ treated mouse CD8^+^ T cells (20). Consistent with these previous reports, TCR-induced Myc mRNA (Figure 2D) and protein levels (Figure 2E, Supplemental Figure 1) were substantially impaired by TGFβ in OT-I T cells. By contrast, levels of *Max* mRNA, that encodes the Myc binding partner Myc-associated factor X (MAX) were marginally increased by TGFβ whereas levels of *Mxd1* and *Mxd4*, encoding transcriptional repressor Max dimer (MAD) proteins, were substantially elevated (Figure 2D).

In T cells, Myc expression is predominantly regulated by a combination of TCR- and IL-2-dependent signals (21). Consistent with previous publications (12, 20) TGFβ reduced OVA-N4 and OVA-T4-induced OT-I T cell IL-2 production (Supplemental Figure 1A). We sought to determine whether a paucity of autocrine IL-2 production accounted for, or contributed to, reduced Myc expression in TGFβ-treated OT-I T cells. *Il2* transcription peaks within the first 6-12h, then drops substantially, following T cell activation (22, 23). We reasoned that delaying the addition of TGFβ for 24h would uncouple indirect effects of reduced *Il2* levels on Myc expression. When added after 24h of TCR-stimulation, TGFβ suppressed Myc expression, as assessed at 48h, albeit to a lesser extent than when added from timepoint 0h (Supplemental Figure 1B). To determine whether exogenous IL-2 could rescue Myc expression, OT-I T cells were stimulated with either OVA-N4 or OVA-T4 ± TGFβ ± IL-2. Data indicated that IL-2 replenishment had no effect on the levels of Myc expression in T cells stimulated with OVA-N4 and TGFβ, whereas it was slightly but not significantly increased in OVA-T4 stimulated T cells (Supplemental Figure 1C).

Previous studies have linked TGFβ-mediated suppression of NK cell function to inhibition of mTOR (24). By contrast, levels of TCR-induced phosphorylation of the mTORC1 downstream target ribosomal protein S6 were unimpeded by TGFβ in OT-I T cells (Figure 2F). Together these analyses indicate that TGFβ-mediated suppression of CD8^+^ T cell activation is linked to repression of Myc activity and expression, independent of effects on IL-2 production and mTOR activity.

### TGFβ impairs amino acid uptake and protein synthesis

Myc transcriptional activity is required for TCR-induced T cell metabolic reprogramming. In particular, Myc-dependent T cell activation is strongly associated with the regulation of amino acid (AA) transporter expression and AA uptake (3, 25). Similarly, our RNA-Seq data indicated that TGFβ treatment resulted in decreased gene expression of several amino acid transporters including *Slc1a5* (encoding Alanine/Serine/Cysteine-preferring transporter 2 [ASCT2]), *Slc3a2* and *Slc7a5* (encoding CD98/ Large neutral amino acid transporter 1 [LAT1]) (Figure 3A). Furthermore, FACS analysis confirmed that TCR-induced upregulation of cell surface ASCT2 (Figure 3B) and CD98 (Figure 3C) was impeded by TGFβ. Kynurenine, a metabolite of tryptophan, can be detected by virtue of its spectral fluorescent properties to assess amino acid transport via SLC7A5/CD98 in T cells by flow cytometry (26). Consistent with decreased transcript and protein levels of CD98, uptake of kynurenine was reduced in TGFβ-treated OT-I T cells (Figure 3D), suggesting that TCR-induced upregulation of AA uptake was impaired. A fundamental role for amino acids in cell biology is to act as building blocks for protein synthesis. Levels of nascent protein synthesis in activated T cells were assessed using incorporation of O-propargyl-puromycin (OPP) and flow cytometry. Importantly, TGFβ treatment substantially impaired TCR stimulated upregulation of OPP incorporation by OT-I T cells (Figure 3, E-F).

**Figure 3:**
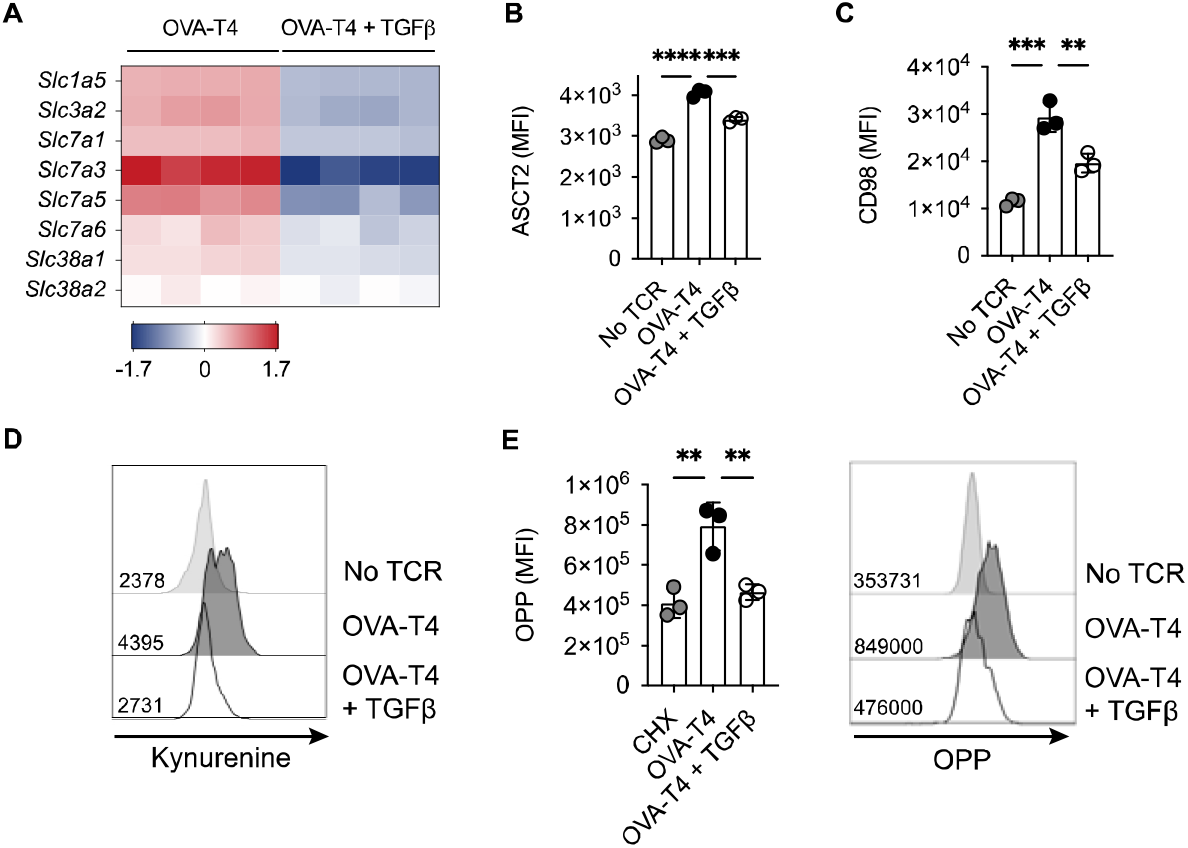
TGFβ impairs amino acid uptake and protein synthesis. OT-I TCR transgenic T cells were stimulated with SIITFEKL (OVA-T4) peptide in the presence or absence of TGFβ for 24 **(A)** or 48 **(B-E)** hours. **(A)** The heatmap shows log_2_ fold changes in expression of selected amino acid transporter-encoding genes as determined by RNA-sequencing. Levels of surface expression of ASCT2 **(B)** and CD98 **(C)** and uptake of kynurenine (D) were determined by flow cytometry. Incorporation of O-propargyl-puromycin (OPP) was detected by Click chemistry staining and flow cytometry (E). For flow cytometry analyses data shown represent mean fluorescence intensity (MFI) values. Individual data points represent biological replicates **(A)** or technical replicates **(B-E)** from 1 of 3 repeated experiments. ** p<0.01, *** p<0.001, **** p<0.0001 as assessed by 1-way ANOVA, with Tukey’s multiple comparisons test.

### TGFβ inhibits TCR-induced upregulation of glycolytic metabolism

Myc expression also mediates the upregulation of glycolytic metabolism following TCR stimulation (25). Analysis of RNA-Seq data indicated that TGFβ modestly impaired mRNA expression of the majority of glycolytic enzymes, with the exception of *Hk1, Aldoc* and *Ldh2b* (Figure 4A). Furthermore, levels of *Slc2a1* and *Slc2a3* (encoding glucose transporters 1 and 3 (GLUT1/3), respectively) were significantly lower in OT-I T cells activated with OVA-T4 + TGFβ as compared to OVA-T4 alone. Consistent with RNA-Seq data, FACS analysis demonstrated that TCR-induced surface expression of GLUT1 protein was impaired in TGFβ-treated cells (Figure 4B). T cell uptake of the fluorescent glucose analogue 2-(*N*-(7-Nitrobenz-2-oxa-1,3-diazol-4-yl)Amino)-2-Deoxyglucose (2-NBDG) is not dependent upon GLUT1 or GLUT3 expression in activated T cells (27), but was also substantially impaired by TGFβ (Figure 4C). Furthermore, analysis of lactate secretion (Figure 4D) and extracellular acidification rate (ECAR) (Figure 4, E-G) determined that TGFβ substantially impairs TCR-induced upregulation of glycolytic metabolism.

**Figure 4:**
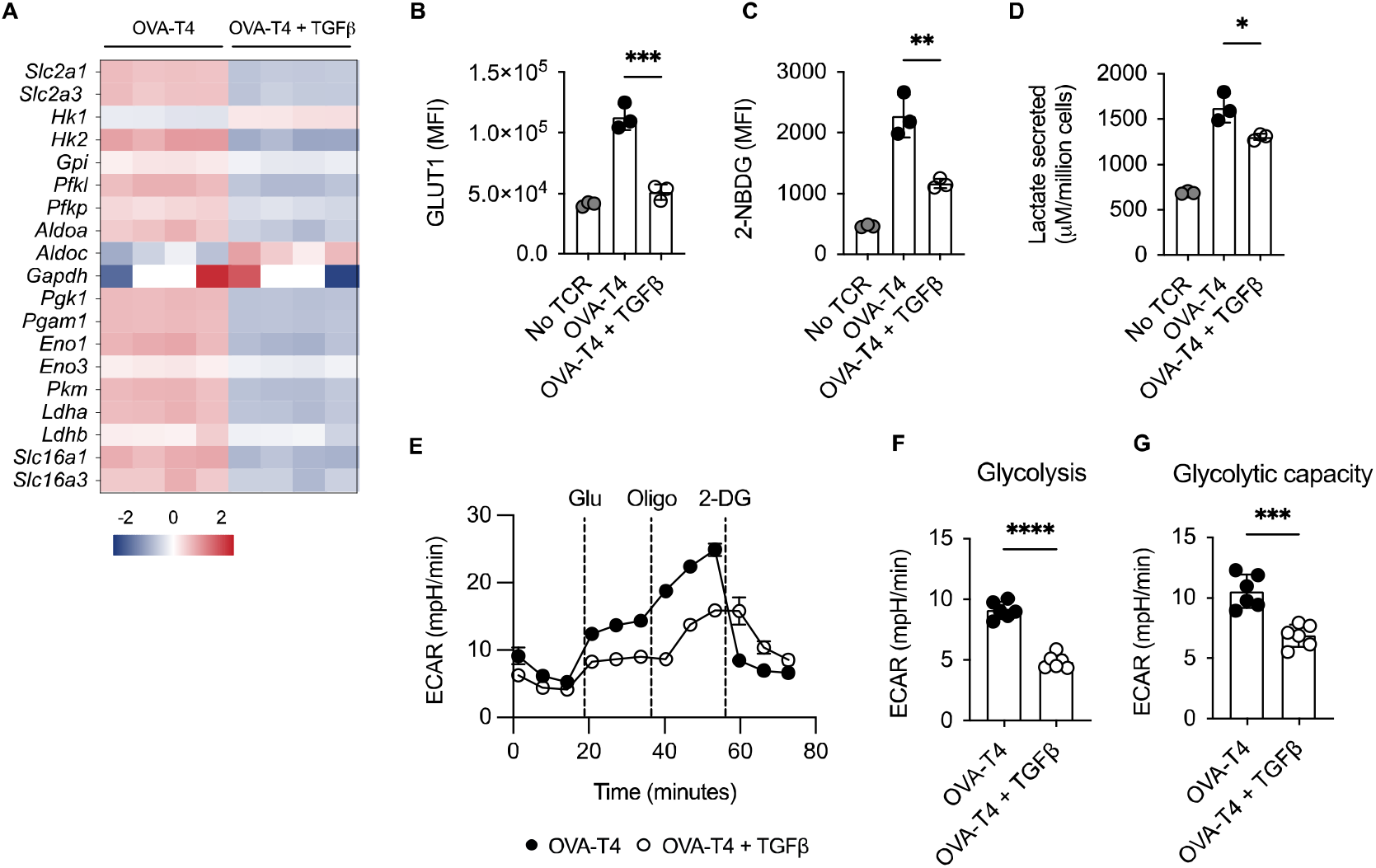
TGFβ impairs CD8^+^ T cell glycolytic metabolism. OT-I TCR transgenic T cells were stimulated with SIITFEKL (OVA-T4) peptide in the presence or absence of TGFβ for 24 **(A, D)** or 48 **(B, C, E-G)** hours. **(A)** The heatmap shows log_2_ fold changes in expression of glycolytic enzyme-encoding genes as determined by RNA-sequencing. Levels of surface expression of glucose transporter (GLUT)1 (B) and uptake of 2- (*N*-)7-nitrobenz-2-oxa-1,3-diazol-4-yl)Amino)-2-deoxyglucose (2-NBDG) (C) were determined by flow cytometry. For flow cytometry analyses data shown represent mean fluorescence intensity (MFI) values. Individual data points represent technical replicates (n=3) from 1 of 3 repeated experiments. (D) Lactate secretion was determined by analysis of deproteinated culture supernatants using a luminescent assay. Individual data points represent biological replicates (n=3) from 1 of 4 repeated experiments. (E) Seahorse metabolic analysis determined extracellular acidification rates (ECAR) of OT-I T cells. Dotted vertical lines represent injection times for glucose (Glu), oligomycin (Oligo) and 2-deoxyglucose (2-DG) in the Glycolytic Stress Test™. Levels of Glycolysis (F) were calculated by subtracting background ECAR values from those determined following glucose injection, whilst glycolytic capacity (G) represent ECAR values following oligomycin injection. For Seahorse metabolic analyses, error bars represent SD and individual data points represent technical replicates (n=6) from 1 of 4 repeated experiments. NS – not significant, * p<0.05, ** p<0.01, *** p<0.001, **** p<0.0001 as assessed by 1-way ANOVA, with Tukey’s multiple comparisons test (B-D) or Student’s *t*-test (F, G).

Finally, we assessed the impact of TGFβ on T cell mitochondrial metabolism by performing a Mitostress Test using the Seahorse analyser. Analysis of oxygen consumption rates determined (Figure 5A) that both basal respiration (Figure 5B) and spare respiratory capacity (Figure 5C) were comparable in TCR as compared to TCR+TGFβ treated OT-I T cells. Furthermore, flow cytometry analysis of tetramethylrhodamine, methyl ester (TMRM) (Figure 5D) and MitoTracker Green (Figure 5E) fluorescence in OT-I T cells indicated that TGFβ did not impinge upon T cell mitochondrial membrane potential or mitochondrial mass, respectively.

**Figure 5:**
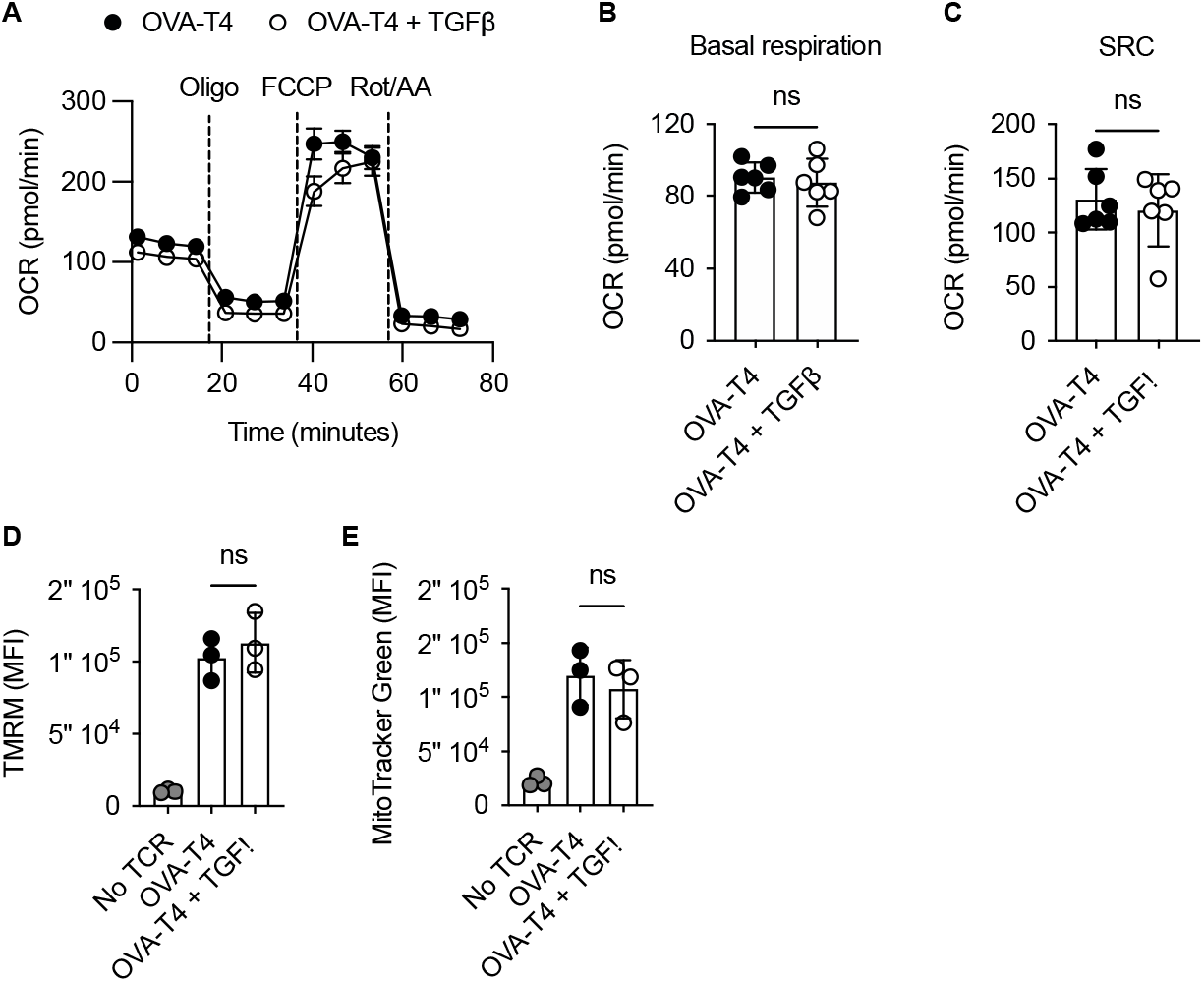
TGFβ inhibits CD8 T cell activation independently of effects on mitochondrial metabolism. OT-I TCR transgenic T cells were stimulated with SIITFEKL (OVA-T4) peptide in the presence or absence of TGFβ for 48 hours. (A) Seahorse metabolic analysis determined oxygen consumption rates (OCR) of OT-I T cells. Dotted vertical lines represent injection times for oligomycin (Oligo), carbonyl cyanide-p-trifluoromethoxyphenylhydrazone (FCCP) and rotenone/antimycin A (Rot/AA) in the MitoStress Test™. Basal respiration **(B)** and spare respiratory capacity (SRC) **(C)** were determined as baseline OCR values and OCR values following FCCP injection, respectively. Mitochondrial membrane potential (D) and mitochondrial mass (E) were assessed by flow cytometry analysis of tetramethylrhodamine methyl ester (TMRM) and mitotracker green fluorescence, respectively. For flow cytometry analyses, data shown represent mean fluorescence intensity (MFI) values. Individual data points represent technical replicates (n=3-6) from 1 of 3 repeated experiments. NS – not significant as assessed by Student’s *t*-test **(A-C)** or 1-way ANOVA, with Tukey’s multiple comparisons test **(D, E).**

## Discussion

CD8^+^ T cell activation depends on transcriptional and metabolic reprogramming mediated by Myc (1). Here we have shown that an important mechanism by which TGFβ impedes early T cell activation is through repression of Myc expression upon TCR-priming. Consequently, T cells activated in the presence of TGFβ display deficient engagement of Myc downstream signalling that is reflected in reduced upregulation of glycolytic and AA-dependent metabolism. These metabolic defects are likely to contribute towards the disruption of TCR-induced CD8^+^ T cell activation, proliferation and effector function.

TGFβ is a master regulator of effector and regulatory T cell activity during autoimmune reactions. Loss of TGFβ signalling in genetically modified murine models leads to aberrant hyperactivation of T cells against self-peptides (28–30). In accordance with our data, these previous studies indicate that TGFβ has a particularly important role in limiting T cell responses upon weak activation of TCR signalling (31, 32). Beyond its implications in autoimmunity here we have shown, in experiments using ID8 ovarian carcinoma cells, that cancer-cell derived TGFβ also restricts T cell activity more efficiently in response to low affinity as compared to strong agonist peptides. In the context of anti-cancer immunity, these results add to a body of work indicating that TGFβ neutralisation could boost responses against tumour associated antigens leading to an increased tumour immunogenicity. A limitation of immune checkpoint blockade therapy is that weakly immunogenic (or “cold”) tumours respond poorly to treatment. Anti-TGFβ therapies might therefore be a promising strategy to convert “cold” tumours into “hot” tumours and improve immune checkpoint blockade when administered in combination.

TGFβ suppresses cytotoxic functions of CD8^+^ T cells through well-defined direct interactions of SMAD proteins with *Gzmb, Ifng* and other inflammatory gene loci, which has been described as a mechanism of immune escape during anti-tumour responses and has also been observed in this study (20). In particular, our transcriptomic data shows that *Gzmb* is the most downregulated gene in the presence of TGFβ highlighting its powerful inhibition of effector functions. Furthermore, we also observed substantial repression of *Eomes* gene expression. Eomesodermin is a key transcription factor involved in CD8^+^ T cell differentiation, particularly through its cooperation with Tbet to promote memory formation. Whether TGFβ exposure during TCR-priming primarily results in deficient CD8^+^ T cell activation or whether it shapes T cell fate and differentiation is likely to be highly context-dependent. Indeed, in the context of an inflammatory cytokine milieu, TGFβ has a well-established role in driving the differentiation of inflammatory Tc17 cells, whilst KEGG pathway analysis of our RNA-Seq data confirmed an upregulation of Tc17/Th17-associated genes in TGFβ-treated cells.

Our RNA-seq data has allowed us to gain a broader understanding of how TGFβ shapes the T cell transcriptome and to dissect possible underlying mechanisms by which it suppresses early T cell activation. After performing KEGG and ChEA analysis we found that Myc-regulated genes were strongly enriched in the TGFβ-downregulated gene set. Myc regulates metabolic reprogramming induced upon TCR-priming of naïve cells to promote the exit from quiescence. Specifically, Myc orchestrates the upregulation of genes involved in glucose and AA transport, mainly *Slc2a1* and *Slc7a5*, in addition to metabolic enzymes related to glycolysis and glutaminolysis. Loss of Myc expression in CD8^+^ T cells prevents TCR-induced blastogenesis, proliferation and differentiation, indicating the essential role of Myc in the initiation of T cell responses (1, 3, 33). Our results confirmed that the marked *Myc* repression observed in the RNA-Seq data results in lower Myc protein levels as well as an important suppression of its main downstream metabolic pathways. Interestingly, while we observe an impairment of glycolysis, protein synthesis and nutrient transport, no significant differences were shown when mitochondrial metabolism was assessed. These results suggest that the effects of TGFβ on T cell metabolism during early stages of activation are a specific consequence of impaired Myc expression rather than being an indirect effect of an overall reduced T cell activation state. In contrast to the current work suggesting minimal effects of TGFβ on mouse CD8^+^ T cell mitochondrial metabolism, previous studies interrogating human CD4^+^ T cells indicated that TGFβ signalling influences respiration via direct effects of SMAD proteins within mitochondria (8). Further work is required to determine if these divergent effects of TGFβ signalling on T cell metabolism are explained by species differences, T cell activation status or intrinsic differences between CD4^+^ and CD8^+^ T cells.

An association between TGFβ signalling and Myc has been previously described in keratinocytes, oligodendrocytes, NK and T cells. Indeed, some studies determined that SMAD proteins are able to directly interact with *Myc* promoter in transformed cells (34). An investigation by Gu et al. showed that Smad4 can also act independently of TGFβ signalling inducing Myc-dependent proliferation in T cells (35). Consequently, Smad4-deficient T cells fail to upregulate Myc expression leading to a deficient expansion. Whether the activation of TGFβ signalling recruits Smad4 and prevents the Smad4-dependent stimulation of Myc expression is still unknown. Further investigations are required to elucidate how TGFβ signalling controls Myc expression in CD8^+^ T cells, which may identify potential mechanisms to prevent the TGFβ-mediated suppression of T cell activation.

A surprising finding in our study has been the lack of effect of TGFβ on mTOR activity considering the important role of AA levels in mTOR regulation (36–38). It is worthy of note that TCR-stimulated naïve T cells are fully competent to induce mTOR activation prior to upregulation of AA transporters. Our data indicate that whilst TGFβ may limit AA uptake, this is insufficient to compromise mTOR activation over a short time-span. Nonetheless, previous studies performed in NK cells and in both CD4^+^ and CD8^+^ T cells have suggested that, in some circumstances, mTOR activity decreases upon TGFβ treatment (8–11). Interestingly, Zaiatz-Bittencourt and colleagues described that mTOR activity in human NK cells was not affected with short TGFβ treatments and that inhibitory effects could only be observed when treatment was sustained for longer periods (5 days) (10). These results suggest that, in specific contexts, the impact of TGFβ signalling on mTOR activity might not be direct and that the effects may only be detected upon the accumulation of other events that could subsequently impact mTOR function.

In summary, we have described here that TGFβ signalling significantly interferes with the Myc-induced transcriptional signature that occurs during early stages of CD8^+^ T cell activation, leading to deficient metabolic reprogramming. Therefore, TGFβ signals during CD8^+^ T cell activation mediate direct inhibition of cytotoxic gene expression by SMAD proteins and metabolic disruption following disruption of Myc expression, both of which contribute to defective CD8^+^ T cell activation. These inhibitory effects are likely important for the maintenance of self-tolerance *in vivo* but may also contribute to T cell immunosuppression during anti-cancer immune responses.

## Methods

### Mice

All experiments were performed using OT-I *Rag1^-/-^* or OT-I mice. OT-I *Rag1^-/-^* mice were maintained at the University of Leeds St. James’s Biomedical Services (SBS). OT-I mice were maintained at the animal facility of University of Lausanne. All experiments were performed using age- and sex-matched mice (7-12 weeks).

### Cell lines

ID8 ovarian carcinoma cells expressing ovalbumin variants (ID8-OVA-N4 and ID8-OVA-T4) were originally a gift from Prof. Dietmar Zehn (Technical University of Munich, Germany). ID8-OVA-N4 and ID8-OVA-T4 cells were maintained in IMDM supplemented with 5% FBS, 50μM β-mercaptoethanol, penicillin-streptomycin and ciprofloxacin (0.5%).

### Cell culture

OT-I T cells were obtained from lymph nodes or spleens of OT-I or OT-I *Rag1^-/-^* transgenic mice. OT-I T lymphocytes were cultured in Iscove’s Modified Dulbecco’s Medium (IMDM, Gibco) supplemented with 5% FBS (Gibco), penicillin-streptomycin (Gibco) and β-mercaptoethanol (Gibco, 50μM). OT-I T cells were activated with 10^-8^ SIINFEKL or SIITFEKL (Cambridge Peptides) in the presence or absence of recombinant human TGFβ (Peprotech, 5ng/ml). Where indicated, OT-I T cells were maintained inactivated *in vitro* (‘No TCR’ condition) with recombinant mouse IL-7 (Peprotech, 10ng/ml). For IL-2 replenishment experiments, recombinant human IL-2 (Peprotech, 1ng/ml) was added to the wells at timepoint 0h. For IL-2 secretion experiments, OT-I T cells were cultured in the presence of blocking CD25 mAb (Biolegend). After 24h of stimulation, supernatants were collected and IL-2 concentration was determined using the Mouse DuoSet IL-2 ELISA (R&D Systems), following manufacturer’s instructions. For ID8-OT-I co-culture experiments, either ID8-OVA-N4 or ID8-OVA-T4 cells were seeded in 48-well plates at 2×10^4^ cells/well. After 4h of incubation to allow adhesion, freshly isolated OT-I T cells were added into the wells (2×10^5^ cells/well) and cultured for 48h. Brefeldin A (2.5mg/ml) was added during the last 6h of culture for optimal detection of cytokine expression by flow cytometry.

### Flow Cytometry

The following conjugated antibodies were used: CD8β - PECy7 (clone YTS156.7.7, 1:400, BioLegend), CD25 – PE (clone 3C7, 1:200, BioLegend), CD44 – APC Cy7 (clone IM7, 1:200, BioLegend), CD71 – FITC (clone RI7217, 1:200, BioLegend), CD98 – PercP Cy5.5 (clone RL388, 1:200, BioLegend), IFNγ – AF488 (clone XMG1.2, 1:100, BioLegend), Granzyme B – BV421 (clone GB11, 1:100, BioLegend), c-Myc – APC (clone D84C12, 1:50, Cell Signalling Technologies), GLUT1 – AF488 (clone EPR3915, 1:400, Abcam). ASCT2 (clone V501, 1:50, Cell Signalling Technologies) and pS6 (clone D68F8, 1:200, Cell Signalling Technologies) were stained with unconjugated antibodies. An additional staining was performed with secondary goat Anti-Rabbit IgG (1:1000, Life Technologies). For assessment of intracellular markers, cells were fixed and permeabilised using FoxP3 fix/perm buffer (eBioscience). For live/dead discrimination, LD Aqua or Zombie NIR were used (LifeTechnologies and BioLegend, respectively). For assessment of protein synthesis, cells were treated with O-propargyl-puromycin (OPP, 20μM, Jena Bioscience) for 30’. OPP was detected using Click-iT Plus AF488 picolyl azide kit (Life Technologies). Cycloheximide (CHX, 100μg/ml, SigmaAldrich) was added as a negative control 15’ prior to OPP treatment. For nutrient uptake assays, T cells were treated with either the fluorescent glucose analogue 2- (*N*-(7-Nitrobenz-2-oxa-1,3-diazol-4-yl)Amino)-2-Deoxyglucose (2-NBDG, 50μM, 1h, Abcam) or kynurenine (800μM in HBSS buffer, 4min, SigmaAldrich), respectively (39).

### Seahorse Analysis

For ECAR and OCR measurements, OT-I T cells were plated in a Cell-Tak coated (22.4μg/ml) Seahorse XFe96 Microplate (10^5^ cells/well). For MitoStress tests, Seahorse assay media was supplemented with glucose (10mM), pyruvate (1mM) and glutamine (2mM). OCR values were acquired with Seahorse XFe96 Analyser at basal levels and upon sequential injection of oligomycin (1μM), carbonyl cyanide-p-trifluoromethoxyphenylhradone (FCCP, 1.5μM) and rotenone / Antimycin A (0.5μM), following standard Seahorse XFe96 protocol. For Glycolysis Stress Test, Seahorse media was supplemented with pyruvate (1mM) and glutamine (2mM), and ECAR values were acquired at basal levels and after sequential injection of glucose (10mM), oligomycin (1μM) and 2-deoxyglucose (50mM), following standard Seahorse XFe96 protocol.

### Lactate Assay

To measure lactate secretion, 10^6^ OT-I T cells were stimulated *in vitro* with OVA-T4 ± TGFβ in 48-well plates. After 24h, levels of lactate in deproteinised culture supernatants were assessed following manufacturer’s instructions (Kit 700510, Cayman Chemicals). Four biological replicates per condition were included.

### RNA extraction and RNA-Seq

OT-I T cells (5×10^6^ cells/sample) were stimulated with OVA-T4 ± TGFβ, as described above. After 24h, cells were lysed with TRI Reagent^®^ (Zymo Research, R2050-1-50) and RNA was extracted using Direct-zol™ RNA Miniprep Plus (Zymo Research, R2070), following manufacturers’ instructions. RNA purity was checked using Nano Drop™ spectrophotometer (A_260_/A_280_ and A_260_/A_230_ > 1.8). RNA-seq and data analysis was performed by Novogene Bioinformatic Technology Co. (Hong Kong). For Kyoto Encyclopaedia of Genes and Genomes (KEGG) pathway analysis and ChIP Enrichment Analysis (ChEA), RNA-Seq data was processed using Enrichr.

### Statistics

Statistical significance (p-value < 0.05) was determined performing Student’s t-test, Mann-Whitney test, and one- or two-way ANOVA with Tukey’s multiple comparison tests using GraphPad Prism, as described in figure legends. Dots in graphs represent replicate samples and error bars represent SDs, unless otherwise stated. For RNA-Seq data, statistical significance was determined using DESeq2 package and p-values were adjusted with the Benjamini-Hochberg procedure.

### Study Approval

Mouse breeding and experiments performed at University of Leeds (UoL) were reviewed by the UoL Animal Welfare and Ethical Review Committee (AWERC) and were approved by and subject to the conditions of a UK Home Office Project Licence (PDAD2D507) held by RJS.

## Author Contributions

HCH and RJS designed the study. HCH performed the majority of experiments. HCH, GP, PG and RJS acquired and analysed the data. NV and GPC supervised parts of the project. HCH and RJS wrote the manuscript. All authors read and discussed manuscript drafts.

## Acknowledgements

This work was supported by grant 23269 from Cancer Research UK (to RJS) and a University of Leeds PhD scholarship (to HCH). NV and HCH are supported by grant KFS-4993-02-2020-R from the Swiss Cancer League.

