## Supplemental Data for "TGFβ limits Myc-dependent TCR-induced metabolic reprogramming in CD8^+^ T cells"

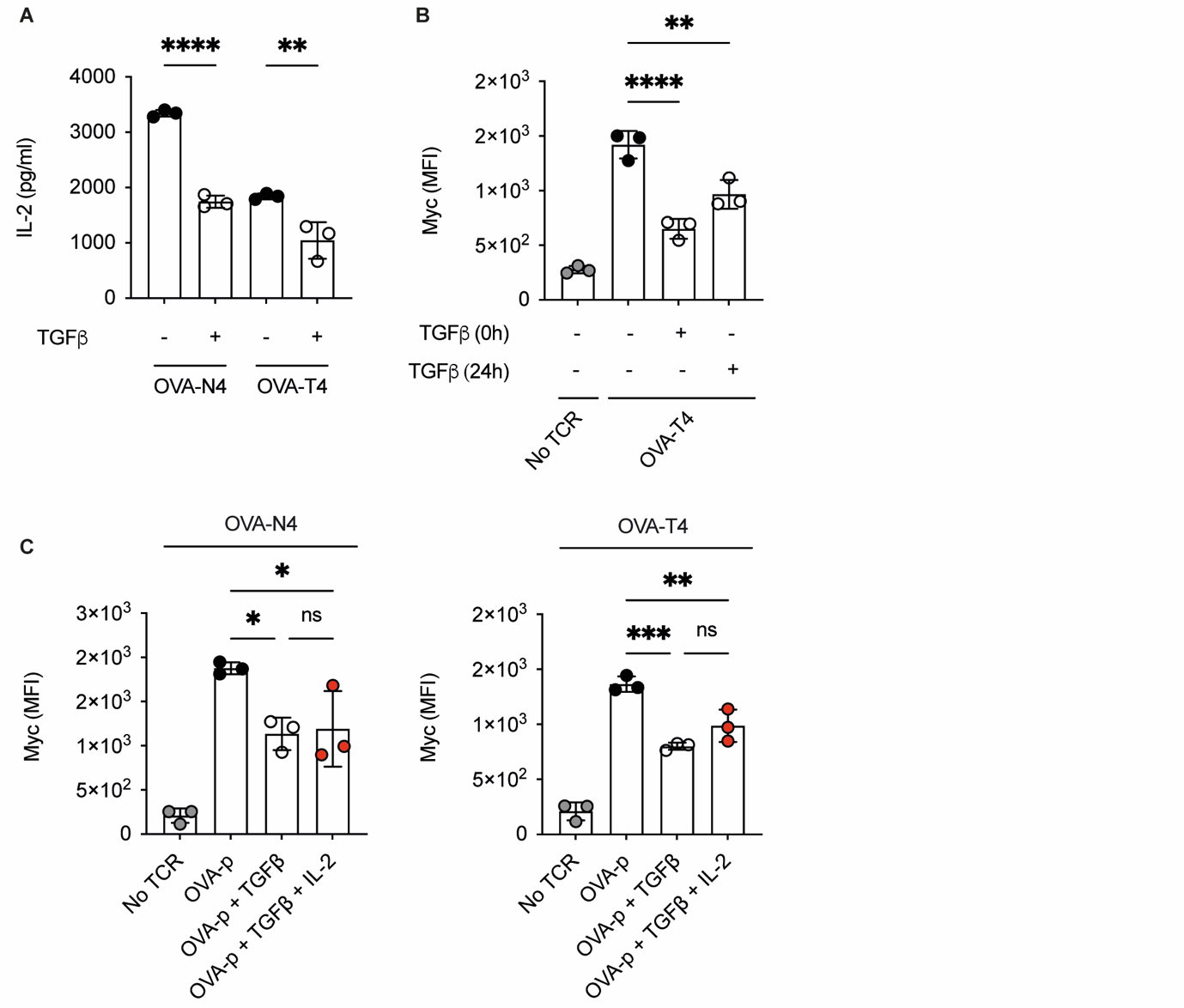

**Supplemental Figure 1. TGFβ-mediated Myc repression is independent of IL-2 depletion.**

OT-I T cells were stimulated with SIINFEKL (OVA-N4) or SIITFEKL (OVA-T4) in the presence or absence of TGFβ for 24h (**A**) or 48h (**B, C**). (**A**) OT-I T cells were stimulated in the presence of CD25 blocking mAb to prevent IL-2 consumption. IL-2 levels in supernatants were determined by ELISA. To assess the role of IL-2 depletion on the TGFβ-mediated repression of Myc expression, OT-I T cells were treated with TGFβ only after 24h of TCR-priming (**B**) or together with hIL-2 (1ng/ml) from timepoint 0h (**B**). Individual data points represent technical replicates from 1 of 3 repeated experiments. ns – not significant, * p<0.05, ** p<0.01, *** p<0.001, **** p<0.0001 as assessed by 1-way ANOVA, with Tukey’s multiple comparisons test.

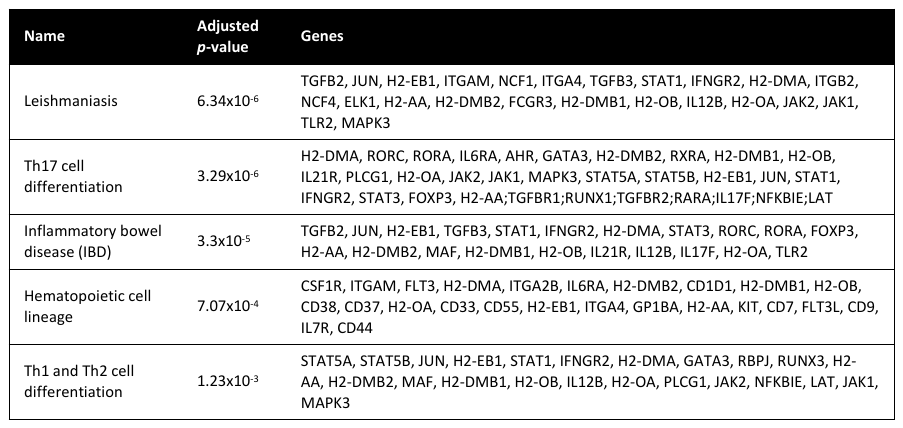

**Supplemental Table 1. KEGG pathway analysis of upregulated genes by TGFβ.**

List of top 5 pathways upregulated by TGFβ in RNA-Seq dataset of OT-I T cells stimulated with SIITFEKL (OVA-T4) ± TGFβ for 24h, as determined by KEGG analysis using the platform Enrichr. Table includes list of genes identified as differentially expressed (*p* adjust < 0.05, fold-change > 1.5) in RNA-Seq dataset.
